# SAKPE: A Site Attention Kinetic Parameters Prediction Method for Enzyme Engineering

**DOI:** 10.1101/2025.04.30.651216

**Authors:** Jia-He Qiu, Zongying Lin, Ke-Wei Chen, Tian-Yu Sun, Xian Zhang, Li Yuan, Yonghong Tian, Yun-Dong Wu

## Abstract

The quantitative determination of enzyme kinetic parameters traditionally relies on experimental methods that are both time-intensive and costly. Machine learning models have demonstrated significant potential for predicting enzyme kinetic parameters in recent years. Despite this promise, these methods face challenges, including limited training data, inadequate sensitivity to subtle mutations, and poor alignment with practical enzyme engineering contexts. Here, we introduce SAKPE (Site Attention Kinetic Parameters Prediction Method for Enzyme Engineering), a novel machine-learning framework designed to predict enzyme kinetic parameters with enhanced accuracy in practical application scenarios. By incorporating protein representation, substrate representation, and protein representation with weights for important sites, SAKPE significantly outperforms existing methods in predicting enzymatic kinetic parameters, including turnover number (*k*_cat_), Michaelis constant (*K*_m_), and inhibition constant (*K*_i_). Incorporating protein representation with weights for important sites enables SAKPE to effectively capture the impact of mutations, especially mutations of important sites and their surrounding amino acids of interest in enzyme engineering, on enzyme kinetics parameters. SAKPE offers a robust and practical tool for predicting enzyme kinetic parameters, providing a superior tool for enzyme engineering scenarios such as enzyme design, directed evolution, and industrial applications.

## 1. Introduction

As natural biocatalysts, enzymes play a central role in living organisms by facilitating the transformation of simple molecules and complex compounds by accelerating chemical reactions^1,2^. With advances in gene cloning and recombinant DNA technology, enzymes can now be synthesized efficiently via microbial expression systems, offering powerful catalytic tools for producing high-value chemicals and pharmaceuticals^3,4^. Developing enzyme systems optimized for industrial applications requires systematic screening and engineering to enhance catalytic activity and stability^5,6^. The catalytic efficiency and substrate affinity of enzymes are commonly evaluated using the Michaelis-Menten kinetic model^7^. Traditional experimental approaches, such as environmental screening^8,9^, metagenomic mining^10^, high-throughput screening of directed enzyme evolution^11^, and other methods, have proven effective in enzyme discovery and optimization. However, the explosive growth of high-throughput genomics and proteomics has led to an exponential expansion in the number of known enzyme sequences. For instance, the UniProt database now contains over 250 million protein sequences, including more than 40 million sequences of enzymes. Nevertheless, fewer than 0.3% of protein sequences have functional annotations^12^. This stark imbalance highlights the limitations of current experimental methodologies in scaling to match the growing sequence space, posing significant challenges to the comprehensive characterization of natural enzymes^13^.

With the rapid progress in computational biology, more cost-effective and efficient enzyme computational screening has gradually become a pivotal means to replace traditional experimental screening^14–17^. Among them, predicting enzyme kinetic parameters is particularly critical. By directly predicting properties, such as enzyme catalytic efficiency, efficient enzymes suitable for chemical synthesis can be identified^18–26^. For instance, David and his colleagues integrated proteomics data under various growth conditions with computational flux predictions to calculate the effective turnover rates for intracellular processes catalyzed by the corresponding macromolecules (*k*_eff_) under diverse conditions^27^. Similarly, Heckmann et al. successfully employed machine learning techniques to predict the *k*_cat_ value of enzymes for *Escherichia coli* using average metabolic flux and other features as part of a diverse set^28^. Following these studies, the DLKcat model utilizes the convolutional neural network (CNN) to extract enzyme sequence features and the graph neural network (GNN) to capture substrate properties, thereby establishing the first general model for predicting the turnover number (*k*_cat_) of enzymes^29^. Moreover, the TurNup model, using the advanced protein language model ESM-1b, was trained on a meticulously curated dataset and demonstrated notably robust performance in out-of-distribution tests, even with a relatively small dataset size. It is also generalizable to enzyme sequences with less than 40% sequence identity^30^. Concurrently, the UniKP model leverages the pre-trained language model ProtT5-XL-UniRef50 to encode enzyme information and incorporates two key environmental factors, pH and temperature, to achieve precise predictions of enzyme kinetic parameters^31^. Numerous other models have also introduced innovative approaches to predicting kinetic parameters from various perspectives, offering fresh insights into the field^32–38^. More importantly, these computational methods can be integrated with techniques such as ancestral sequence reconstruction (ASR)^39^, greedy accumulated strategy for protein engineering (GRAPE)^40^, binding free energy calculations^41^, and other emerging computational approaches. To a certain extent, it provides optional tools to meet the demand for customized properties of enzymes with both high activity and high stability^19,42–44^.

Although enzyme computational screening has made significant progress to a certain extent, existing computational methods still face many challenges. First, the limited size of the training dataset and the incomplete diversity of enzyme and substrate types mean that the model may learn insufficient information^42^. Many existing machine learning models have overfitting problems and limited generalization capabilities^45^. In addition, most existing enzyme kinetics parameters prediction models do not identify the important sites in the enzyme sequence, making it difficult to assess the impact of mutations on enzyme activity sensitively^45^. The lack of this important information led to the enzyme kinetic parameter prediction tools being constrained in the following key practical application scenarios of enzyme engineering. The catalytic activity of an enzyme is closely linked to its three-dimensional structure, particularly the precise configuration of its catalytic site or binding pocket. This structural specificity allows the enzyme to selectively recognize and bind its substrate, thereby enabling efficient catalysis of chemical reactions^46,47^. Reetz et al. found that beneficial mutations that enhance enzyme catalytic chiral selectivity are primarily concentrated in the substrate binding pocket, thus creating the combinatorial active-site saturation test (CAST)^48^, iterative saturation mutagenesis (ISM)^49^, and focused rational iterative site-specific mutagenesis (FRISM)^50^. This semi-rational design method effectively narrows the range of mutation sites from which to select. Directed evolution of substrate binding pockets has become an effective and widely used strategy in enzyme engineering^51–53^. In enzyme engineering, rationally designed experiments favor mutations closer to the catalytic pocket (including mutations at important sites)^54^. In contrast, the existing general enzyme kinetics prediction parameters models do not expand the features for the changes before and after the key amino acid sites of the mutants change, resulting in insensitivity to changes in enzyme kinetic parameters.

To make up for the shortcomings of existing methods, we proposed a new machine learning framework, SAKPE (Site Attention Kinetic Parameters Prediction Method for Enzyme Engineering), which aims to improve the performance of enzyme kinetic parameter prediction, especially to improve the prediction ability of the model for mutants with mutations at important sites. SAKPE is based on a common dataset containing extensive *k*_cat_, *K*_m_, and *K*_i_ data and incorporates more comprehensive feature information, including amino acid sequences, SMILES^55^ strings of substrates, and important sites of enzymes (such as catalytic sites, binding sites, and other important sites). By providing the model with information about important sites of the enzyme, SAKPE is endowed with the ability to learn the impact of these sites on catalytic activity, thereby accurately predicting the kinetic parameters of the enzyme, especially when mutations of important sites and their surrounding amino acids occur.

We hope that SAKPE can provide more accurate computational support for the design and modification of enzymes and promote the application of enzyme engineering technology in industry and biotechnology.

## 2. Results

### 2.1. Model architecture and dataset of SAKPE

The SAKPE framework employs a hierarchical architecture combining multi-modal feature extraction with the gradient boosting model (Figure 1). First, the dataset used by SAKPE is a comprehensive dataset constructed based on the two mainstream enzyme kinetic parameter databases, BRENDA^56^ and SABIO-RK^57^, after careful extraction and integration (Figure 1a, Supplementary Figure S1). To ensure a rigorous evaluation, random partitioning was implemented to create the training dataset (90%) and the test dataset (10%), while maintaining similar distributions of enzyme classes and parameter ranges between the splits (see the “Dataset Acquisition and Processing” section for details). Subsequently, in the representation extraction stage, multiple pre-trained protein language models and small molecule language models were used to extract features and generate representations for the SMILES^55^ (Simplified Molecular Input Line Entry System) strings of the input enzyme protein sequence and substrate. Regarding protein features, the ESM-C^58^ protein language model was used to map the amino acid sequence into a 1152-dimensional representation. The global embedding representation of the entire protein sequence was obtained through the mean pooling strategy. At the same time, to enhance the perception of important sites of the model, the site importance of the enzyme protein sequence was further encoded to form a site weight representation, which was combined with the protein representation, to extract the protein representation with weights for important sites. Regarding substrate features, the SMILES strings of substrates were input into the dedicated molecules pre-trained model Mole-BERT^59^ for encoding. Then, a 300-dimensional embedding representation was generated for each character, which ultimately constituted the overall representation of the substrate (Figure 1b). Finally, in the training stage, the protein sequence representation, site importance representation, and substrate representation were jointly input into the subsequent gradient boosting model module for training to achieve the prediction task of enzyme kinetic parameters (Figure 1c).

**Fig. 1:**
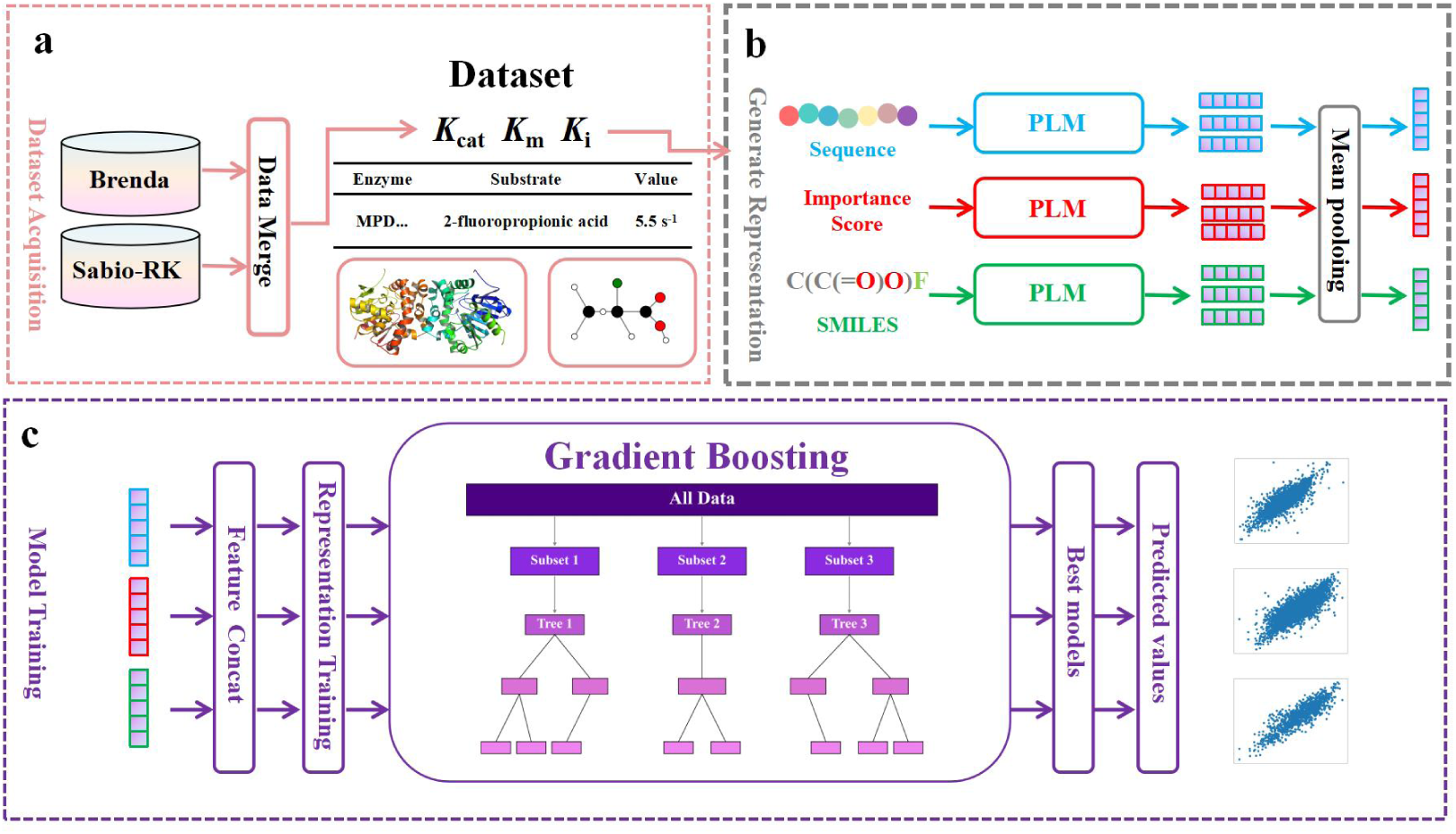
Overall of the SAKPE. **a** Construction, integration, and partitioning of the dataset based on the BRENDA and SABIO-RK databases. **b** Extraction of embedded representations from protein sequences and substrate using the pre-trained language model, incorporating protein representation with weights for important sites. **c** Input the protein and substrate representations into the gradient boosting model to predict enzyme kinetic parameters.

A systematic analysis of the dataset was conducted, and a critical analysis revealed various notable characteristics of the dataset (Figure 2). The SAKPE dataset contains a total of 31,507 *k*_cat_, 53,310 *K*_m_, and 10,841 *K*_i_ entries across seven major enzyme classes (EC 1-7). First, by comparing the distribution of *k*_cat_, *K*_m_, and *K*_i_ in the entire data, wild-type samples, and mutant samples, it was shown that when higher *k*_cat_ and lower *K*_m_ and *K*_i_ were used as the optimal standard, the wild-type enzyme generally performed better than the mutant in all three parameters (Figure 2a). This result may be attributed to the fact that during the natural selection process, the enzyme retains a large number of functionally conserved sites, which are often used as targets for functional verification or engineering mutations. Therefore, these mutations usually hurt the catalytic efficiency of enzymes^60^. Subsequently, the mutation sites of the mutants were analyzed, and it was found that a considerable number of mutations involved amino acids related to important sites. In the *k*_cat_ dataset, approximately 28% of the mutants harboured mutations at important sites. In the *K*_m_ and *K*_i_ datasets, these proportions were 30% and 37%, respectively (Figure 2b). Overall, mutants accounted for 35%, 32%, and 15% in the *k*_cat_, *K*_m_, and *K*_i_ datasets, respectively, which is highly representative and shows that mutants are an essential part of enzyme engineering (Supplementary Figure S2a). In addition, a larger proportion of mutations were concentrated in important sites and their surrounding amino acids, suggesting that these sites are also often considered to contribute more to changes in enzyme performance (Supplementary Figure S2b). Furthermore, the distribution ratio of the seven major enzyme entries (EC 1-7) in the dataset was analyzed. The results showed that oxidoreductases (EC 1), transferases (EC 2), and hydrolases (EC 3) accounted for more than 75% of the total number of entries. The distribution ratio of mutants in different enzyme types was shown simultaneously (Figure 2c). Finally, the distribution characteristics of the kinetic parameters for the seven types of enzymes were analyzed, and it was found that there were significant differences in the distribution of parameters such as *k*_cat_, *K*_m_, and *K*_i_ for each enzyme. Notably, the top 15 substrates and the top 15 organisms with the highest frequency in the dataset were counted, and the enzyme kinetic parameters corresponding to different substrates or host sources showed significant differences (Supplementary Figure S3). These distribution differences emphasize the diversity of the dataset and also pose a higher challenge to the generalisation ability of the model.

**Fig. 2:**
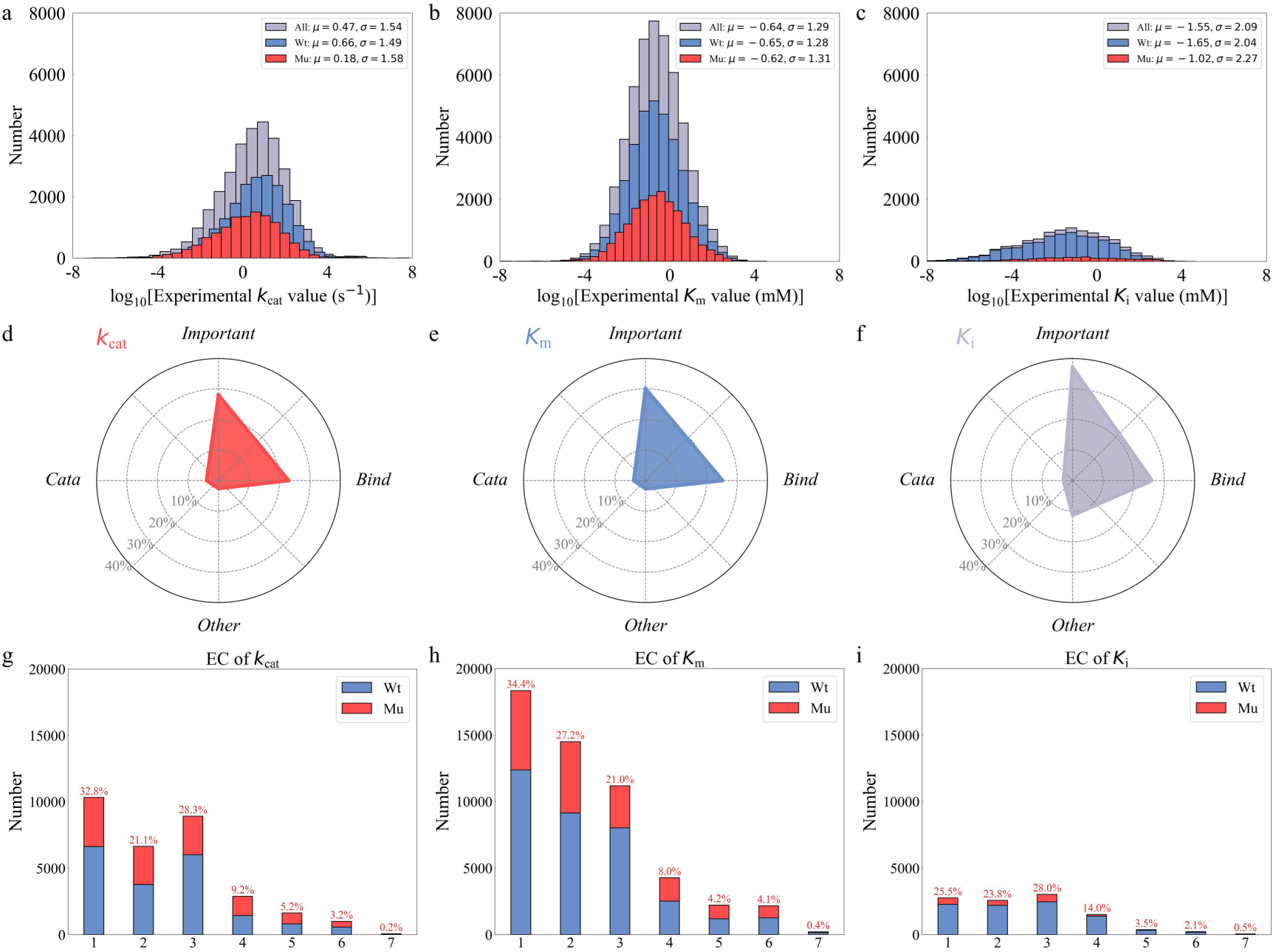
Composition and Statistical Feature Analysis of the SAKPE Dataset. **a-c** Comparison of the overall data distributions for wild-type and mutant samples for the three parameters *k*_cat_ (a), *K*_m_ (b), and *K*_i_ (c). **d-f** Statistical overview of important site-related mutations in the mutant dataset, including the proportions of catalytic sites, binding sites, and other important site-related mutations for the three parameters *k*_cat_ (g), *K*_m_ (h), and *K*_i_ (i). **g-i** Analysis of the proportion of the seven major enzyme classes (EC primary classification: EC 1: Oxidoreductase; EC 2: Transferase; EC 3: Hydrolase; EC 4: Lyase; EC 5: Isomerase; EC 6: Ligase; EC 7: Translocase) corresponding to the *k*_cat_ (d), *K*_m_ (e), and *K*_i_ (f) parameters.

### 2.2. Performance of SAKPE in the test dataset

The pre-trained models employed were systematically evaluated to generate representations of protein sequences and substrates, covering a variety of representation methods. For protein representation, a variety of embedding methods based on protein language models, including ESM-C, ESM2^61^, and ProtT5-XL-UniRef50 (ProtT5)^62^, were used, and they are widely employed in the field of protein structure and function prediction. For substrate representation, not only were traditional rule-based molecular fingerprint methods, such as Morgan and MACCSKeys^63^ examined, but also several embedding representations generated based on chemical molecular language models, including Uni-Mol^64^, Uni-Mol2^65^, and Mole-BERT, were evaluated. These models can provide diverse perspectives on the representation of protein sequences and substrates, thereby enabling a comprehensive evaluation of the applicability of different representation methods in the downstream task of predicting enzyme kinetic parameters.

Model performance was rigorously evaluated using four established metrics: coefficient of determination (R^2^), Pearson correlation coefficient (PCC), root mean square error (RMSE), and mean absolute error (MAE) (see the “Performance Evaluation” section for details). Our analysis revealed that the ESM-C + Mole-BERT combination achieved superior performance in kcat prediction (R^2^ = 0.719, RMSE = 0.799), with statistically significant advantages across all metrics (Figure 3a, d). For predicting *K*_m_ and *K*_i_ tasks, while showing a slight disadvantage (ΔR^2^ = 0.01), the ESM-C + Mole-BERT combination maintained comparable accuracy (Figure 3b-c, e-f). Crucially, all metrics exhibited strong concordance, as evidenced by the consistent ranking patterns with the PCC value of 0.848 and the MAE value of 0.555. This shows that the choice of representation method has a universal impact on model performance, and there is a strong consistency between different tasks (Supplementary Figure S4). Based on the balanced performance of ESM-C and Mole-BERT in different prediction tasks, the subsequent comparison and in-depth analysis in this article will mainly focus on this combination to explore its prediction performance in various task contexts.

**Fig. 3:**
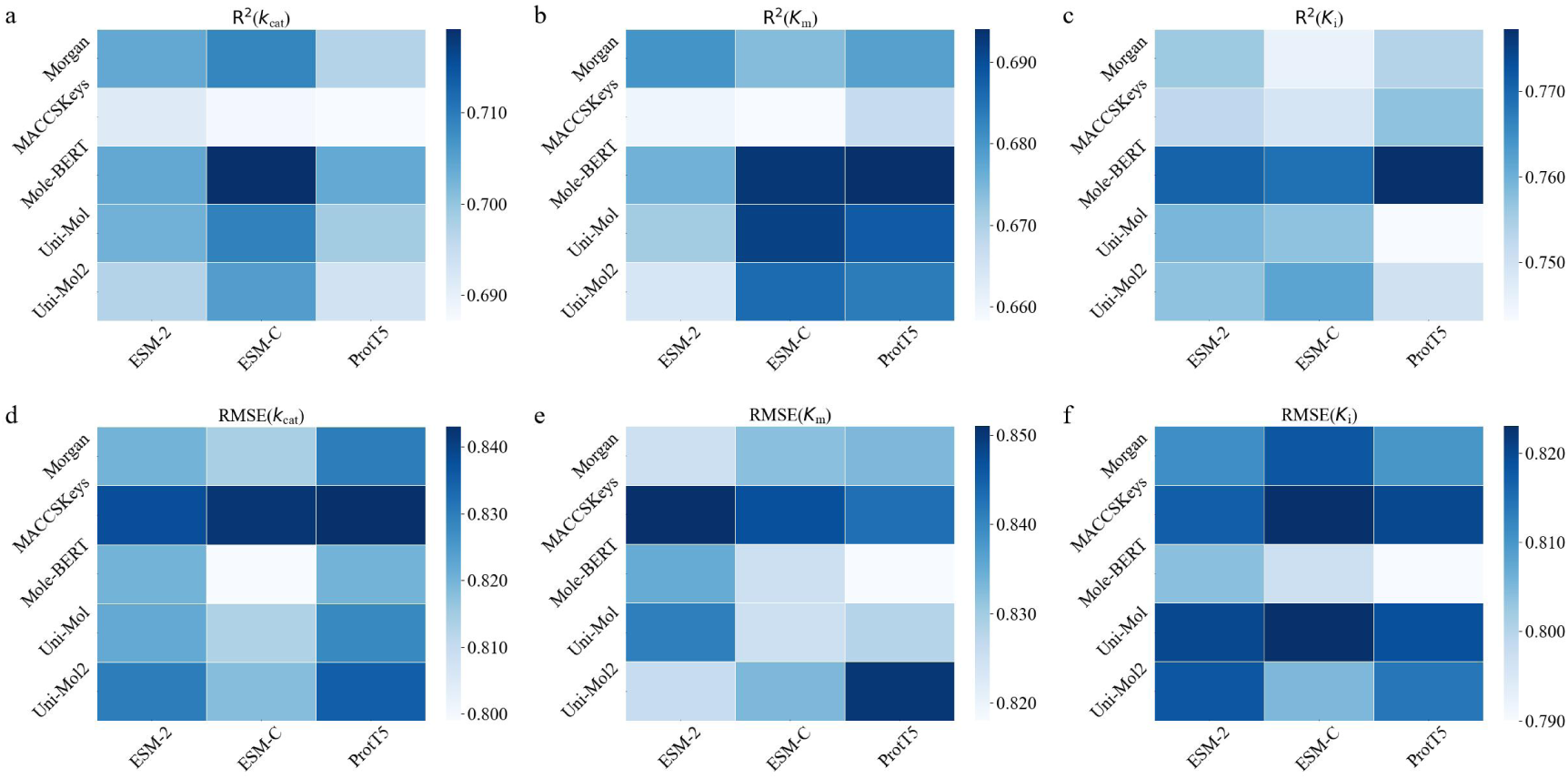
Heatmap Depicting the Performance of Various Representation Method Combinations in Enzyme Kinetic Parameter Prediction. **a-c** Heatmaps illustrating the R^2^ for the prediction tasks of *k*_cat_ (a), *K*_m_ (b), and *K*_i_ (c), respectively. **d-f** Heatmaps depicting the PCC for the *k*_cat_ (d), *K*_m_ (e), and *K*_i_ (f), prediction tasks, respectively.

Each small square in the heatmap corresponds to a specific combination of a protein pre-training language model (on the horizontal axis) and a small molecule pre-training language model (on the vertical axis). The color scale (blue to white) indicates values from high to low, respectively.

### 2.3. Performance of SAKPE in different subsets of the test dataset

To comprehensively evaluate the applicability and robustness of the model in different tasks, several subsets were divided with representative features from the test dataset, and a systematic performance analysis was carried out. The division of these subsets covers multiple dimensions, including different enzymatic parameter intervals (especially extreme values in the two-tailed distribution), protein sequences with varying numbers of mutation sites, classification of metabolic pathways, enzyme types, substrate types, and organism sources, verifying the great predictive ability of the model from multiple perspectives.

In the prediction tasks for different numerical intervals, it was observed that even in the extreme value area of the enzymatic parameters, the model could still maintain a low RMSE, with all RMSE values not exceeding 1.30. This indicates that SAKPE demonstrated exceptional robustness in predicting edge-case kcat values (Figure 4a). When evaluating the predictive ability of the model for mutants, SAKPE manifests strong sensitivity to mutants (R^2^ = 0.758 ∼ 0.844) even for proteins containing up to five mutations, indicating that the model still had reliable predictive performance in the face of mutation complexity, resilience likely stemming from the site-attention module (Figure 4b). To assess biological relevance, the method proposed by Li et al. for classifying enzymes based on the metabolic pathways they participate in was implemented, and its predictive ability was further explored under different metabolic types^29^. The results showed that the turnover number (*k*_cat_) predicted by SAKPE was significantly higher for enzymes involved in primary metabolism and energy metabolism than for enzymes involved in intermediate metabolism and secondary metabolism. This trend is consistent with existing studies and provides further evidence for the rationality of the prediction results of the model (Figure 4c).

**Fig. 4:**
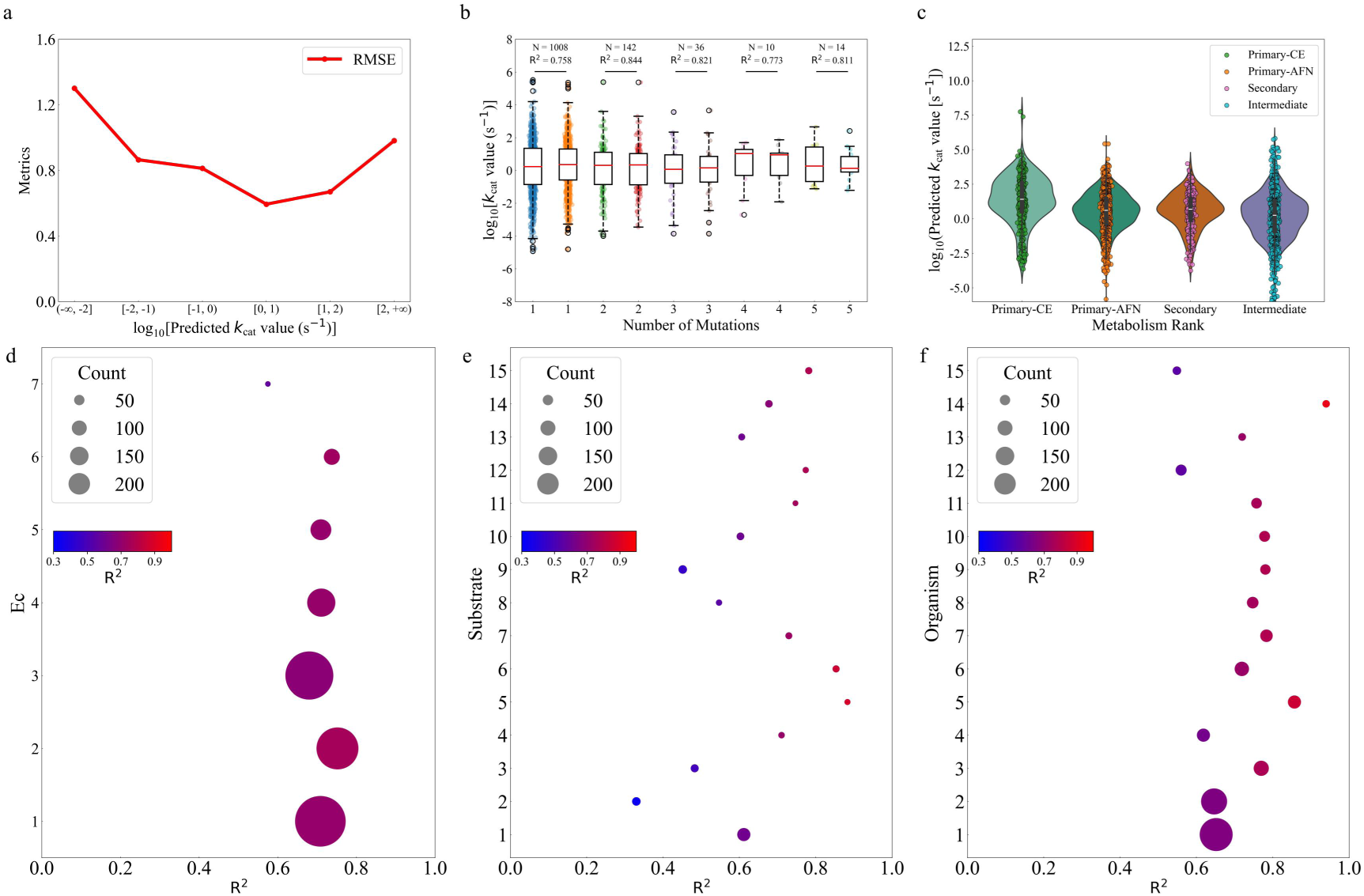
Prediction Performance of SAKPE on Various Subsets. **a** RMSE of predictions evaluated across different ranges of enzymatic parameters. **b** Distribution and R^2^ performance between experimental and predicted values for protein samples grouped by the number of mutations. **c** Distribution of model prediction outcomes for *k*_cat_ of enzymes across different metabolic types. **d-f** Model prediction performance represented by the R^2^ in seven different ec type subsets, the high-frequency substrate subset, and the high-frequency organism source subset, respectively. In the legend, the sphere size is proportional to the number of test cases.

Furthermore, considering that the test dataset covers a variety of enzyme types and that different enzymes have significant *k*_cat_ differences (Supplementary Figure S3), comprehensive EC class analysis revealed robust performance across all seven enzyme types (median R^2^ = 0.710), with translocases (EC 7, n = 72) showing reduced but non-trivial predictive power (R^2^ = 0.575) despite representing only 0.2% of training data (Figure 4d), further verifying the broad applicability of the model. The predictive competency of SAKPE in diverse biological contexts was also evaluated. The top 15 substrates and the top 15 organisms with the highest frequency in the dataset were selected, and the qualified subsets were tested. SAKPE maintaining median R^2^ = 0.652 for 15 most frequent substrates (e.g., ATP: R^2^ = 0.612; D-glyceraldehyde 3-phosphate: R^2^ = 0.782) and median R^2^ = 0.725 for 15 common host organisms (e.g., *Homo sapiens*: R^2^ = 0.652; *Escherichia coli*: R^2^ = 0.646), indicating that the model still has outstanding predictive performance when facing host organisms or substrates with diverse sources (Figure 4e-f).

### 2.4. Comparison with previous models

To establish rigorous benchmarking of enzyme kinetic parameter capabilities, a comprehensive comparison was conducted against two recently published generalizable frameworks: DLKcat and UniKP. Adhering to the original implementation protocols, both models on our curated dataset, using identical data splits and preprocessing pipelines, were retrained to ensure an equitable comparison.

Model performance was quantified comprehensively through five complementary metrics: R^2^, PCC, RMSE, MAE, and within one order of magnitude error (p_1mag_) (see the “Performance Evaluation” section for details).

Across five evaluation metrics, SAKPE achieved superior predictive performance, for example, in the *k*_cat_ prediction task (R^2^ = 0.719 vs. 0.468 and 0.648 for baselines, Figure 5a). Similar conclusions also apply to the *K*_m_ and *K*_i_ prediction tasks (Figure 5b-c), further verifying that SAKPE has wide applicability and significant advantages in various enzyme kinetic parameter prediction tasks. To assess generalizability, low-sequence-similarity subsets (with sequence identity thresholds of 99%, 80%, 60%, and 40% compared to the training sequences) were constructed from the test set (see “the “Dataset Acquisition and Processing” section for details). SAKPE maintained robust performance under this stringent evaluation (Figure 5d-f), outperforming DLKcat and UniKP in *k*_cat_ prediction (R^2^ = 0.488 vs. 0.041 and 0.409 for baselines in the subset with a sequence identity threshold of 99%) and showing similar advantages, or at least maintaining the same level of performance, when extended to *K*_m_ and *K*_i_ tasks. This enhanced generalizability stems from our proposed novel method for protein representation with weights for important sites, as detailed in the next Section.

**Fig. 5:**
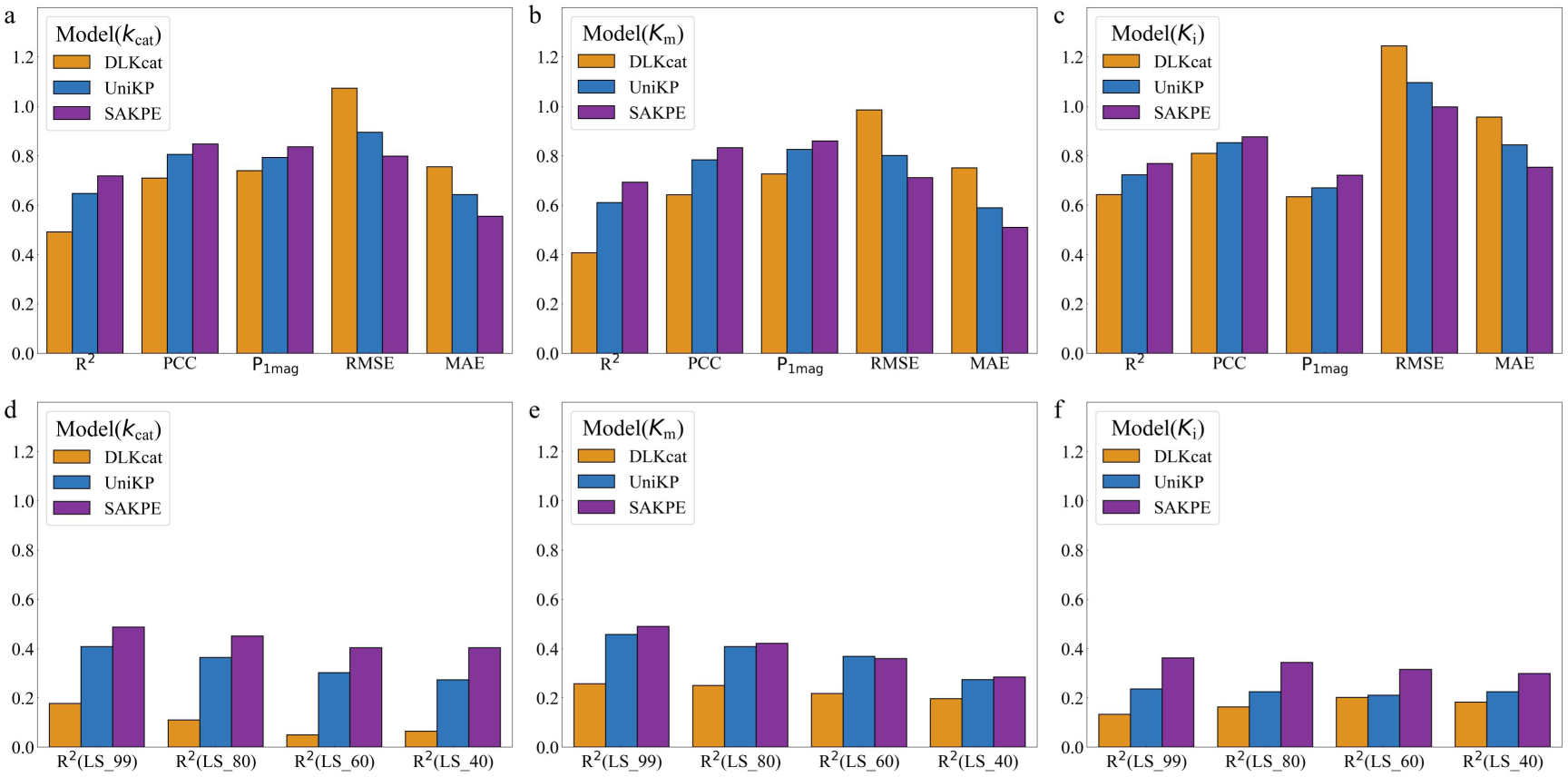
Comparison of the Prediction Performance of DLKcat, UniKP, and SAKPE. **a-c.** Comparison of the evaluation metrics for *k*_cat_ (a), *K*_m_ (b), and *K*_i_ (c) prediction tasks, respectively, on the whole test dataset. **d-f.** Comparison of the R^2^ for the *k*_cat_ (d), *K*_m_ (e), and *K*_i_ (f) prediction tasks, respectively, evaluated on the subset with low sequence similarity. The “LS” denotes low similarity, with the values in parentheses representing the similarity thresholds. A lower threshold value signifies a lower degree of similarity between the test dataset and all sequences in the training dataset.

### 2.5. Significant improvement of the site attention mechanism in practical applications

To quantify the contribution of the site attention module, systematic ablation experiments were performed across multiple test subsets. The whole test dataset analysis revealed consistent improvements when incorporating site importance features, with R^2^ increasing by 0.019 (from 0.700 to 0.719), PCC by 0.011 (from 0.837 to 0.848), and RMSE decreasing by 0.026 (from 0.825 to 0.799) (Figure 6a-b). The generalizability of the model was further evaluated using low-sequence-similarity subsets, with sequence similarity thresholds set from 99 to 10. The SA module significantly enhanced the accuracy when predicting the kinetic parameters of the low-similarity test subset (Figure 6c). Specifically, R^2^ is improved by a range from 0.0686 to 0.122 (Figure 6c). The average number and relative value of the improvement in the prediction results before and after ablation were counted (that is, the average value of the ratio of the absolute value of the predicted difference of each sample before and after ablation to the experimental value) and the average number and relative value of the deteriorated. The results showed that the number (57%) and magnitude (0.30) of improvement were higher than the deterioration in the subset with a sequence identity threshold of 99% (Figure 6d). In addition, SA can also improve the prediction performance of mutants to varying degrees. Specifically, for mutants, performance has been enhanced to a certain degree, mutants not located within 5 Å of important sites exhibited a slight improvement in predictive performance, mutants within 5 Å of important sites showed a significant improvement, while those directly at important sites demonstrated a highly significant enhancement (R^2^ from 0.646 to 0.703) (Figure 6e). Statistical analysis revealed that the relative improvement exceeded that of the deteriorated predictions (Figure 6f). Cross-validation with a stringent low-similarity/mutant intersection subset (n = 6) also confirmed these trends ( Δ R^2^ = +0.705, Supplementary Figure S5). This suggests that our model has great potential for application in enzyme engineering. Furthermore, the ability of the model to identify mutation effects was tested. According to the definition by Kroll^45^, the mutation effect is the difference between the mutant and wild-type parameter changes (minus the mean of both), which shows significant changes before and after ablation^45^, further supporting the positive impact of the site attention module on model performance (Supplementary Figure S6).

**Fig. 6:**
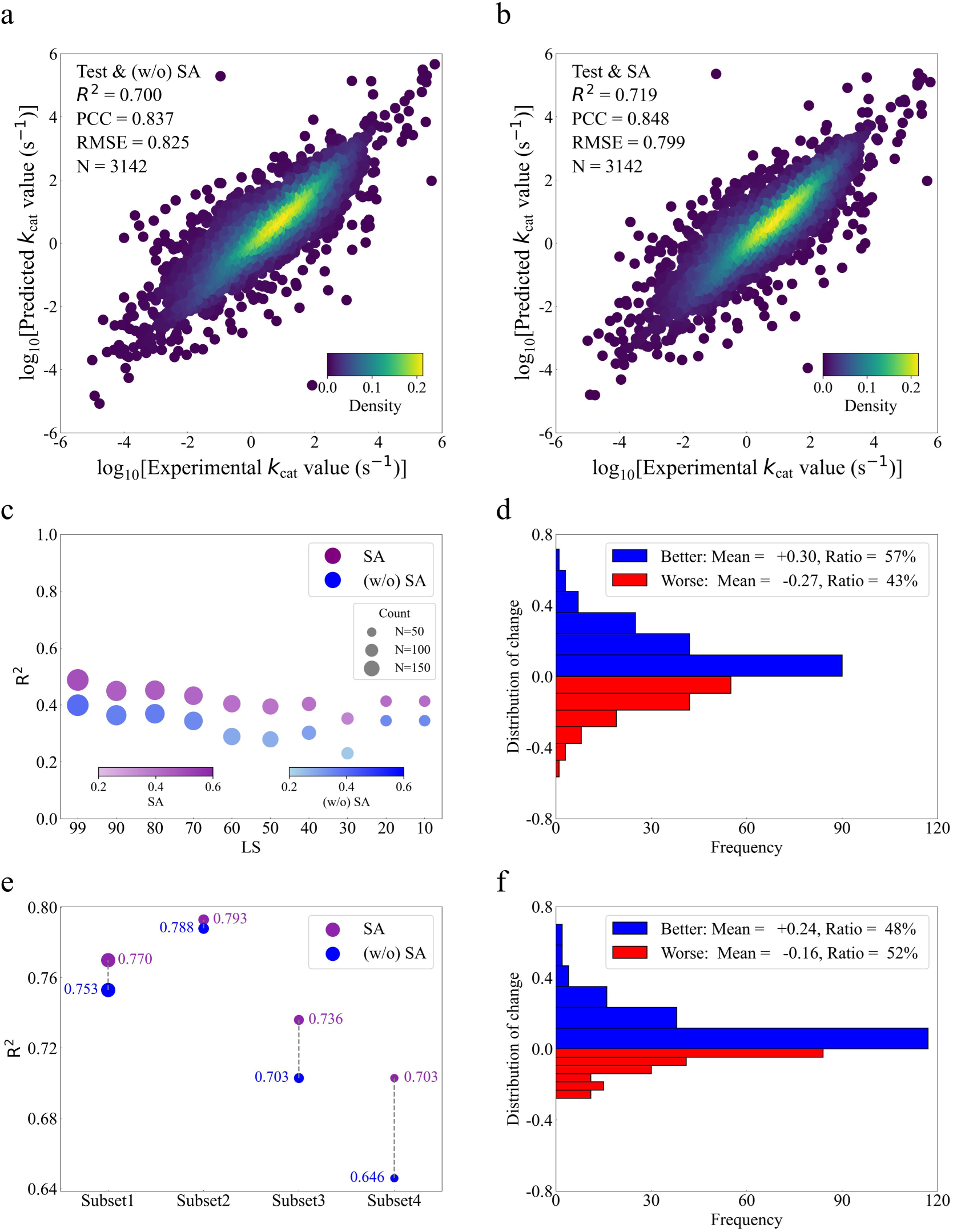
Ablation Test Results of the Site Attention Module. **a,b** Prediction performance of *k*_cat_ values before (a) and following (b) the introduction of protein representation with weights for important sites. Color brightness represents data density. The “SA” denotes site attention. **c** Improvement in performance on the low sequence similarity subset is attributable to the incorporation of protein representation with weights for important sites. As shown in the legend, the size of the sphere is proportional to the number of test datasets. **d** Statistical summary of both the number of improved predictions and the relative improvement observed in the low sequence similarity subset before versus after ablation. The absolute error is the absolute difference between the experimental value and the predicted value. The improvement in absolute error is evaluated by comparing the absolute error differences between the models before and after the module was added. An improvement indicates a reduction in error, while a negative improvement signals an increase in error. To normalize the error improvement, the values are obtained after applying the Sigmoid function. **e** After the introduction of protein representation with weights for important sites, the prediction accuracy of the four kinds of mutant subsets associated with the important site regions was improved to varying degrees. The four subsets are mutants, mutants with mutations not in and around important sites, mutants with mutations in and around important sites, and mutants with mutations in important sites. **f** Statistical summary of both the number of improved predictions and the relative improvement observed in the important site-related mutant subset before and after ablation.

These results show that protein representation with weights for important sites enhances prediction robustness in three key scenarios: novel enzyme sequences outside the training distribution, engineered mutants generated by site-directed mutagenesis, and evaluation of mutation impacts, all of which are vital requirements for practical enzyme engineering applications.

## 3. Discussion

This work proposes a method for predicting enzyme kinetic parameters, SAKPE, that integrates pre-trained models and an important site attention mechanism. A comprehensive dataset was first constructed and used a variety of protein representation methods, including ESM-C, ESM-2, and ProtT5, as well as traditional and chemical molecular language model-based substrate representation methods such as Morgan, MACCSKeys, Uni-Mol, Uni-Mol2, and Mole-BERT, to extract features from the data. Experimental results show that SAKPE has significantly outperformed existing models in predicting parameters such as *k*_cat_, *K*_m_, and *K*_i_. Further ablation tests verify the positive effect of protein representation with weights for important sites. Evaluations on the entire test dataset, low sequence similarity subsets, and important site-related mutant subsets show that the determination coefficient and Pearson correlation coefficient of the model improve after the introduction of important site information. At the same time, the root mean square error and mean absolute error are reduced, showing better generalization ability and prediction stability. Statistical analysis also proved that the prediction improvement after introducing the important site representation was superior in both quantity and magnitude, and enhanced the ability to identify mutation effects. In summary, the SAKPE method effectively captures key factors in enzyme dynamics by integrating pre-trained features with the important site attention module, providing a practical computational tool for screening and designing enzymes in enzyme engineering.

In the extensive evolution of enzymology and the advancement of enzymatic engineering, various theories have been developed to describe how enzymes select and bind to substrates, including Emil Fischer’s lock-and-key principle^66^, Linus Pauling’s hypothesis on transition state stabilization^67^, and conformational selection-based model^68^, among others. These insights make the exploration of mutations in pocket-lining residues a highly rational approach, intimately associated with critical residues and their surrounding regions. This aligns with the fact that the construction of candidate mutant enzymes through saturation mutagenesis of these pocket-lining sites, guided by semi-rational or rational design, has become a widely recognized and effective strategy in enzyme engineering to enhance catalytic activity and stability^69^. Numerous contemporary computational enzyme engineering approaches embody the principle of integrating structural data, active sites in the catalytic pocket, and related biochemical insights to guide rational design^70^. Although the enzyme kinetic parameter prediction tool can predict the mutants of each site mutated to other 19 standard amino acids (i.e. 19×n predictions), and can even be extended to 19^n^ scale parameter predictions, however, given the above-mentioned cognition in enzyme engineering research, enzyme engineering research based on these tools still mainly focuses on mutations of important sites and their surrounding amino acids (Supplementary Table.S1). Based on this, the SAKPE model was developed, which innovatively incorporates important enzyme site information on the basis of providing enzyme sequence and substrate information. This improvement significantly enhances the generalization ability of the model and the prediction accuracy for important sites and their adjacent residue mutations, providing a truly practical and efficient method for the practical application of enzyme kinetic parameter prediction tools in enzyme engineering.

Although the SAKPE model demonstrates robust predictive performance for enzyme kinetic parameters, surpassing existing methods, several challenges remain to be addressed. Firstly, despite including diverse enzyme and substrate types in our training dataset, it still falls short of capturing the full spectrum of enzyme systems encountered in real-world applications. In enzyme engineering, it’s common to test one enzyme on a series of structurally related substrates^71^, yet such data are scarcely documented in existing databases. Secondly, while our model basically achieves state-of-the-art performance on test datasets with low similarity, this limitation persists in practical scenarios. Additionally, enzyme kinetic parameters are highly susceptible to variations in pH and temperature, but these factors are often under-annotated in available datasets. Furthermore, a significant portion of database-derived parameters may not align with the original literature, introducing potential inconsistencies^72^.

Mutations at or near important sites typically result in reduced enzyme kinetic parameters, aligning with the established understanding that important residues, which are generally conserved, are essential for catalytic activity and significantly influence enzyme function. While incorporating important site information markedly enhances the predictive performance of the model, the scarcity of data remains a challenge, as most mutations in the dataset are deleterious. If the dataset could be enriched with information on beneficial mutations, the SAKPE model may achieve superior performance in enzyme discovery and engineering. Furthermore, the kinetic parameters of enzymes do not wholly depend on the important sites of the enzyme and their nearby residues. The remote regulation^73–75^, oligomeric state^76,77^, gates of enzymes^78^, and many other factors^79^ that affect enzyme kinetic parameters were not employed as input features due to the lack of annotation information or annotation tools. Consequently, models that precisely assign weights to enzymes based on their catalytic potential, along with predictive models that leverage prior knowledge for a select group of well-characterized enzymes, offer highly promising avenues for relevant research.

## 4. Materials and Methods

### 4.1. Dataset Acquisition and Processing

The datasets of enzyme kinetic parameters were extracted from the BRENDA^56^ database and the SABIO-RK^57^ database. The datasets of sequences, substrates, and enzyme kinetic parameters were retained. The sequence information of the enzyme was obtained from UniProt^80^, and the entries containing non-standard amino acids were deleted. The SMILES^55^ strings of the substrates were obtained from PubChem^81^. All substrate SMILES strings were standardized using RDKit^63^. It was ensured that the enzyme-substrate pairs were not repeated in the obtained entries, the values were logarithmically transformed, and the overall value distribution was approximately normal. This process generated a total of 31,507 *k*_cat_, 53,310 *K*_m_, and 10,841 *K*_i_ entries, which contained enzyme sequences, SMILES strings of substrates, and enzyme kinetic parameter values. UniProt and EasIFA^82^ annotated the importance of the enzymes, and the weights of important sites were determined. The SAKPE dataset was divided into a training dataset (90%) and a test dataset (10%) using scikit-learn^83^. It was further filtered into subsets according to the sequence identity threshold between the enzyme sequence and the training sequence. The low-identity test dataset was partitioned using mmseqs2^84^. The enzyme sequences in each dataset (*k*_cat_, *K*_m_, and *K*_i_) were clustered using sequence identity thresholds of 10%, 20%, 30%, 40%, 50%, 60%, 70%, 80%, 90% and 99% to obtain the low-sequence identity dataset.

### 4.2. Model Framework

The SAKPE framework was implemented using torch v.2.5.1 + cu 118 and python 3.10. Regarding protein representations, ESM2^61^ is a state-of-the-art protein model trained on a masked language modelling objective. ESMC^58^ focuses on creating representations of the underlying biology of proteins, which scales up data and training compute to deliver dramatic performance improvements over ESM2. ProtT5^62^ is based on the transformer architecture and was pretrained on approximately 45 million non-redundant protein sequences from UniRef50 using a modified masked language modelling. Take ESM-C as an example, each amino acid is converted into a 1152-dimensional vector in the last hidden layer, and the resulting vectors are averaged. The final enzyme representation is a 1152-dimensional vector. The importance of the site was used to obtain a 1152-dimensional protein representation with weights of important sites. Regarding the representations of substrate, Morgan Finger and MACCSKeys Finger were utilized by RDKit to compute these molecular fingerprints. Uni-Mol^64^ is a versatile three-dimensional molecular pre-training framework. Uni-Mol2^65^ is an innovative molecular pre-training model that leverages a two-track transformer to effectively integrate features at the atomic level, graph level, and geometry structure level. Mole-BERT^59^ develops triplet masked contrastive learning to model the varying degree of molecular similarities. For the substrate, the SMILES was generated, and the quality of features extracted by the molecular fingerprints Morgan Finger and MACCSKeys Finger, and the pre-trained Bert-based Mole-BERT and Transformer-based Uni-Mol and Uni-Mol2 were compared. The gradient boosting model XGBoost^85^ was employed for the training stage, which could effectively capture the relationship between the protein representation and substrate, and the enzyme kinetics parameters.

### 4.3. Performance Evaluation

To evaluate the performance of our model, we used five evaluation metrics in the experiment to compare it with existing methods. The coefficient of determination (R^2^) in Equation 1 measures the proportion of variance in the experimental values that is explained by the model prediction. The closer the R^2^ value is to 1, the better the fit between the predicted value and the actual value. The Pearson correlation coefficient (PCC) in Equation 2 quantifies the linear relationship between the predicted value and the experimental measurement. The closer the PCC value is to 1, the stronger the positive correlation is. The root mean square error (RMSE) in Equation 3 and the mean absolute error (MAE) in Equation 4 are used to quantify the difference between the predicted value and the experimental measurement value. The percentage of test predictions that are within one order of magnitude error (p_1mag_) in Equation 5, similar to the definition in CatPred, indicates that a higher value reflects a greater proportion of predicted values falling within one order of magnitude of the experimental measurement value. The assumption is that the logarithms of the four values of the experimental measurement value, computationally predicted value, the average of the experimental measurement value, and computationally predicted value are the average values of the sample numbers, respectively: *y* ^e^, *y* ^p^, *y*^e^, *y*^p^. Their definitions are, respectively, and n is the number of samples:

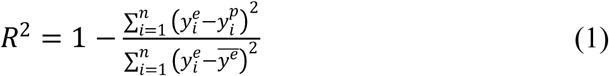

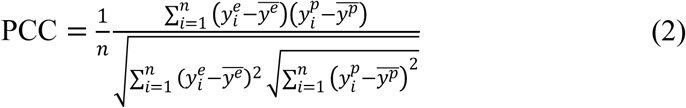

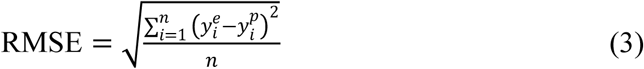

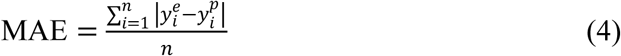

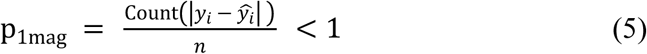

### 4.4. Retraining of Other Baselines

DLKcat and UniKP were selected as baselines, and they were fully reproduced according to the code provided by the authors. Specifically, the training code supplied by the authors was directly used in the original paper and retrained the models of DLKcat and UniKP on our training dataset. The evaluations were performed using a consistent evaluation model. Although DLKcat predicted only *k*_cat_, and UniKP was trained on *k*_cat_ and *K*_m_, both models were used to retrain all three parameters (*k*_cat_, *K*_m_, and *K*_i_) to compare their performance thoroughly.

## 5. Data availability

All data supporting the primary findings of this study are accessible within the manuscript and its Supplementary Information files. The datasets analyzed in this work are publicly available from established databases, including BRENDA^56^(https://www.brenda-enzymes.org/), SABIO-RK^57^ (http://sabio.h-its.org/), UniProt^80^ (https://www.uniprot.org/), PubChem^81^ (https://pubchem.ncbi.nlm.nih.gov/), and KEGG^86^ (https://www.kegg.jp/), as well as supplementary datasets from referenced studies (https://github.com/SysBioChalmers/DLKcat^29^, https://github.com/Luo-SynBioLab/UniKP^31^). Source data are included with this publication. Unless otherwise specified, all data supporting the study’s conclusions are provided in the article, Supplementary Information, or source data files.

## 6. Code availability

The source code for SAKPE is available at https://github.com/HGzyme/SAKPE.

## Acknowledgments

This research was funded by the National Key Research and Development Program of China (No. 2023YFA1506500). The National Natural Science Foundation of China (No. 22403067). This work was supported by Shenzhen Bay Laboratory Supercomputing Center.

## 7. Author contributions

**Jia-He Qiu**: writing - original draft, methodology, formal analysis. **Zongying Lin**: writing - original draft, methodology, formal analysis. **Ke-Wei Chen**: writing - original draft, methodology, formal analysis, supervision. **Tian-Yu Sun**: methodology, formal analysis. **Xian Zhang**: conceptualization, supervision. **Li Yuan**: conceptualization, supervision. **Yonghong Tian**: conceptualization, supervision. **Yun- Dong Wu**: conceptualization, writing - review & editing, supervision.

## Notes

### Competing Interest Statement

The authors have declared no competing interest.

